# Topographic organization of feedback projections to mouse primary visual cortex

**DOI:** 10.1101/2020.07.12.198440

**Authors:** Mai M. Morimoto, Emi Uchishiba, Aman B. Saleem

## Abstract

Context dependent top-down modulation in visual processing has been a topic of wide interest. Recent findings on context dependent modulation, combined with the tools available to investigate network mechanisms in the mouse, make the mouse primary visual cortex an ideal system to investigate context-dependent modulation. However, the distribution of inputs to V1 from across the brain is still relatively unknown. In this study, we investigate inputs to V1 by injecting cholera toxin B subunit (CTB), a retrograde tracer, across the extent of V1. To identify CTB labelled cell bodies and quantify their distribution across various brain regions, we developed a software pipeline that maps each labelled cell body to its corresponding brain region. We found over fourteen brain regions that provided inputs to V1. Higher visual areas (HVAs) provided the most inputs to V1, followed by the retrosplenial, cingulate, and other sensory cortices. As our injections spanned a range of coordinates along the mediolateral axis of V1, we asked if there was any topographic organisation of inputs to V1: do particular areas project preferentially to specific regions of V1. Based on the distribution of inputs from different HVAs, injection sites broadly clustered into two groups, consistent with a retinotopic separation into sites within the central visual field and the peripheral visual field. Furthermore, the number of cells detected in HVAs was correlated to the azimuthal retinotopic location of each injection site. This topographic organization of feedback projections along the medio-lateral axis of V1 suggests that V1 cells representing peripheral vs central visual fields are differentially modulated by HVAs, which may have an ethological relevance for a navigating animal.

## Introduction

Neural activity in the mouse primary visual cortex (V1) is known to be modulated by a variety of contextual signals, including arousal, locomotion, spatial context, spatial attention, or navigation (Niell and Stryker, 2010; Keller et al., 2012; Saleem et al., 2013, 2018; McGinley et al., 2015; Poort et al., 2015; Vinck et al., 2015; Fiser et al., 2016; Jurjut et al., 2017; Pakan et al., 2018; Speed et al., 2020). Perturbations of specific areas have been found to alter some contextual modulations in V1, including mesencephalic locomotor region (MLR) and effects of locomotion (Lee et al., 2014), or anterior cingulate cortex (ACC) and effects of sensorimotor prediction (Leinweber et al., 2017). These studies were limited to investigating the involvement of specific regions projecting to V1. What are the other potential sources of the various contextual signals in V1? The first step towards addressing this question is knowing which regions of the brain project to V1. However, focused effort on mapping brainwide inputs to V1 has been limited. Therefore, in this study we use an unbiased approach using retrograde tracing to characterise cortex-wide inputs to V1.

Higher visual areas are known to provide feedback inputs to V1 and modulate receptive field properties, and activity for factors such as visual features in the surround field (Keller et al., 2020; Nurminen et al., 2018; Vangeneugden et al., 2019). In mouse, over nine discrete cortical areas have been defined as higher visual areas (HVAs) based on architectonic signatures and functional properties (Wang and Burkhalter, 2007; Garrett et al., 2014; Glickfeld and Olsen, 2017; Zhuang et al., 2017). The HVAs have been categorised into two distinct streams based on their anatomical connectivity (Andermann et al., 2011; Marshel et al., 2011; Wang et al., 2011, 2012; Glickfeld and Olsen, 2017; Murakami et al., 2017) and spatial and temporal response properties (Andermann et al., 2011; Marshel et al., 2011; Wang et al., 2011, 2012; Glickfeld and Olsen, 2017; Murakami et al., 2017), proposed to be analogous to the ventral and dorsal streams of primates (Andermann et al., 2011; Marshel et al., 2011; Wang et al., 2011, 2012; Glickfeld and Olsen, 2017; Murakami et al., 2017). Here, we investigated the distribution of inputs from all HVAs to V1.

Topography as a phenomenon has been observed for receptive field position and various tuning properties in V1 and HVAs. Retinotopy across the different HVAs has been shown to be biased: with HVAs medial to V1 generally biased to representing the periphery visual field, while HVAs lateral to V1 biased to the central visual field (Garrett et al., 2014; Zhuang et al., 2017). Growing evidence points to topographic differences in additional functional properties such as binocular disparity tuning, colour tuning, coherent motion processing, and spatial modulation across V1 and HVAs (Aihara et al., 2017; La Chioma et al., 2019; Sit and Goard, 2019). How interconnections between V1 and HVAs contribute to these mesoscale topographies is not well understood. Patterns of connectivity between V1 and HVAs shed light on possible underlying mechanisms. Projection specific calcium imaging studies have revealed reciprocal connections at a single cell level: feedforward projection cells from V1 match tuning properties of recipient HVAs (Glickfeld et al., 2013; Huh et al., 2018), and in turn, V1 cells receive a large portion of their feedback inputs from the HVA they project to (Kim et al., 2020). Anatomical tracing of feedforward projections from V1 to HVAs are known to show striking topography (Wang and Burkhalter, 2007). However, topography of feedback inputs from HVAs to V1 has not been investigated to date.

In this study, we investigated inputs to V1 using the retrograde tracer Cholera Toxin subunit B (CTB). To quantify the inputs to V1 from different brain regions, the first step was to detect labelled cells and identify the brain regions where they are present. As most existing software tools (Fürth et al., 2018; Song et al., 2020; Tyson et al., 2020) require extensive setup and training, we developed a simple and modular software pipeline to quantify labelled cells across the brain. Our pipeline takes 2D images taken with a standard microscope, detects cells, and aligns the images to the Allen CCF, thereby allowing 3D reconstruction of cell positions. Using this software pipeline, we found inputs to V1 originating from visual thalamic nuclei, all cortical HVAs, and over fourteen nonvisual brain regions. The input strength varied across the different areas, and was most prominent from the retrosplenial cortex and HVAs. Inputs from HVAs were organised such that projections from medial HVAs preferentially projected to medial V1 and lateral HVAs projected to lateral V1, resulting in a topographic organization of feedback inputs.

## Results

To investigate inputs to V1, we injected a retrograde tracer, cholera toxin B subunit (CTB) conjugated with Alexa Fluor (488, 555 or 647) across the medio-lateral extent of V1 (Fig 1A). We used retrograde tracing as it offers a direct and simple method of evaluating input projections from across the brain to the injected target region. The tracers were injected bilaterally (7 animals) or unilaterally (2 animals) in 2 or 3 sites within V1, varying along the mediolateral axis to span the extent of V1. To differentiate injection sites within each hemisphere, we used CTB conjugated with different fluorophores (Alexa Fluor 488, 555 or 647). CTB is known to be taken up primarily by axon terminals at the injection site, and retrogradely labels neuronal cell bodies (Nassi et al., 2015). Two weeks post-injection, we sectioned coronal brain slices and stained them with DAPI, and obtained images across the anterior-posterior extent of the brain using standard fluorescent light microscopes (Leica DMi8 or Zeiss Axio Scan). As expected, the retrograde tracer injected into V1 labelled cell-bodies of neurons across various regions of the brain (Figure 1,2).

**Figure 1.**
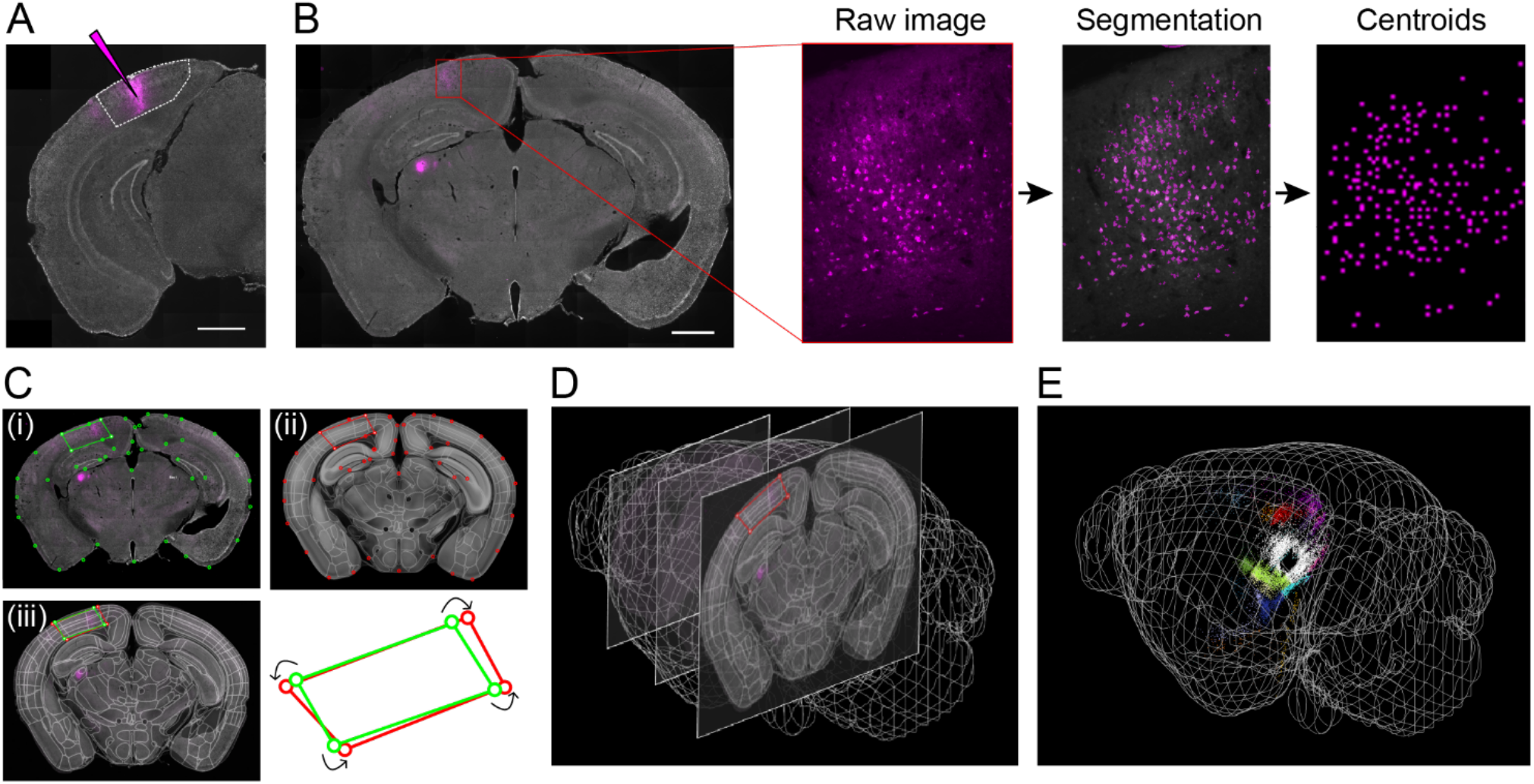
Software pipeline for mapping cell body locations to brain regions through alignment with Allen CCF. **A)** Example injection site of CTB in V1. White dotted line denotes V1 boundary based on the Allen Reference Atlas (ARA). Scale bar: 1 mm **B)** Demonstration of the steps used to detect labelled cells: cells were segmented from the raw image and their centroids were extracted. Scale bar: 1 mm **C)** Images of brain slices were registered to the ARA using SHARP-track. We illustrate an example of this procedure, where transformation points on DAPI image ((i), green) were transformed to fit a Reference slice (from ARA) transformation points ((ii), red). Overlay of (i) and (ii) shown in (iii). **D)** The registration was carried out across all brain slices along the anterior-posterior axis. **E)** An example visualization of cell body locations on the 3D brain model with cells color-coded by area identity.

**Figure 2.**
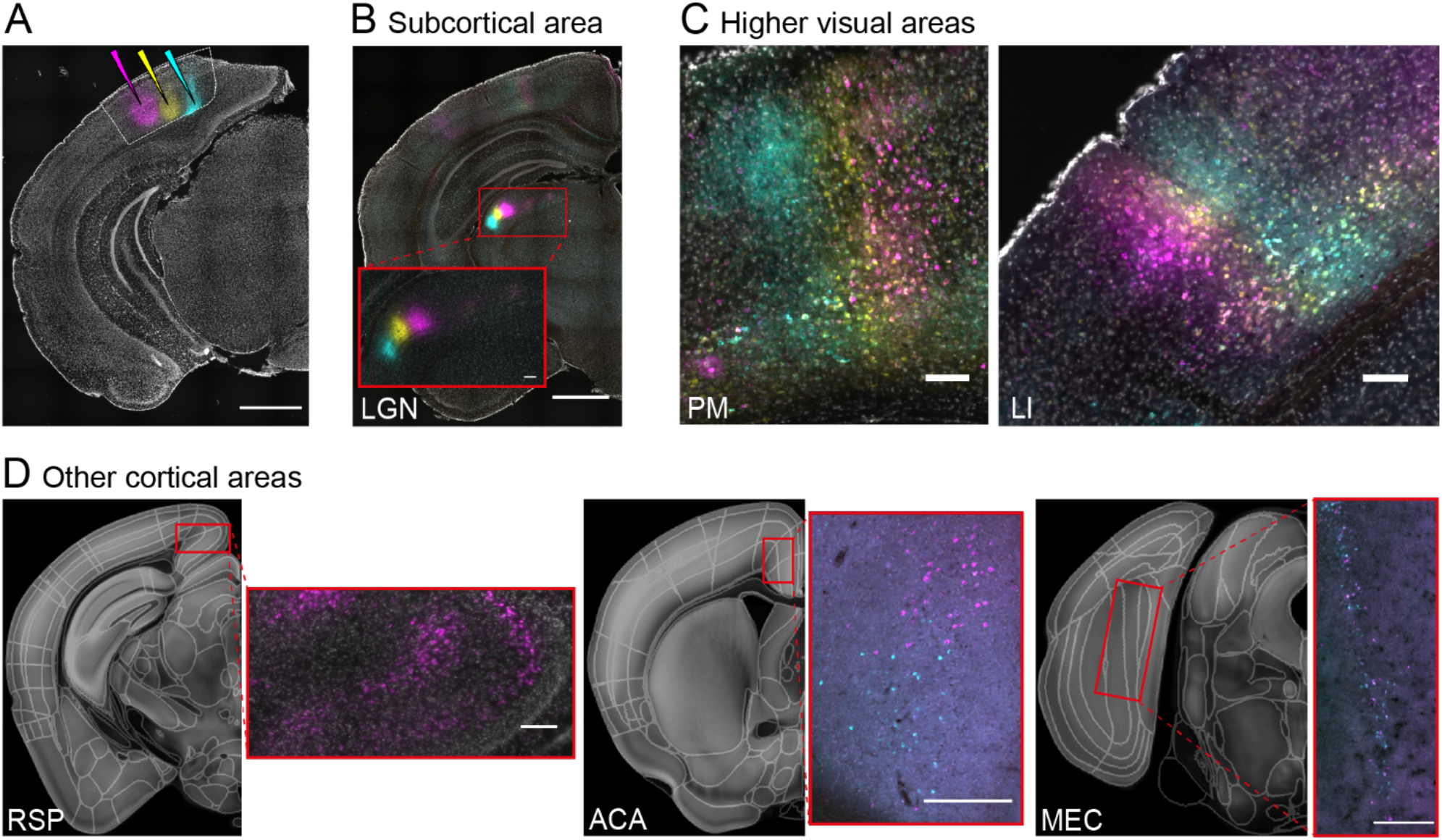
Retrogradely labelled cells detected in various brain regions. **A)** Example of multiple CTB injections in V1 (CTB-488 (cyan), CTB-647 (yellow) and CTB-555 (magenta)). **B-D**) Example areas showing retrogradely labelled cells. Retinotopic organization of cells can be seen in LGN. **(B)** and higher visual areas (PM and LI shown) **(C)**. Cells were observed in other cortical areas such as RSP, ACA, and MEC (For area name abbreviations see Supplementary Table 1) **(D)**. Right side panels correspond to red rectangle regions on the left panel. Scale bars: A-B: 1 mm, C-D: 100 μm.

### Software pipeline for mapping labelled cell bodies to brain regions

In order to identify cell bodies labelled by the tracer, and quantify their occurrence across various brain regions, we developed a software pipeline that maps each labelled cell body to its corresponding brain region, based on the Allen reference atlas (Wang et al., 2020). The first part of the pipeline was finding the centroid locations of the cell-bodies in each brain slice. We achieved this by segmenting CTB labelled cell bodies in the corresponding fluorescent channel image, and extracting centroids from an ellipse fit to these cell bodies (methods, Supplementary Fig. 2). We used the centroid locations to generate a binary mask image, which was one at the centroid locations and zeros elsewhere (centroid mask image, Fig. 1B). The second part of the pipeline was to convert centroid locations to the Allen CCF. For this, we first used Sharptrack (Shamash et al., 2018) to transform our *‘DAPI image’* (DAPI stain) to fit the Allen CCF. Sharp-track allows us to explore the 3D model of the mouse brain (Allen CCF) at different positions and slicing angles, and thus identify the particular slice (*‘reference slice’*) that corresponds to our *DAPI image*. Next, we manually selected corresponding anatomical landmarks in the *DAPI image* and *reference slice*, which were used to locally transform the *DAPI image* to fit the *reference slice* (Fig. 1C). The same image transformations were then applied to the centroid mask image, thus converting the centroid positions into Allen CCF. The final part of the pipeline was to identify the corresponding brain region of each centroid based on the Allen Reference Atlas (defined in Allen CCF), which allowed us to identify the brain area where the centroid of each detected cell was located. Through this semi-automated procedure, we obtained area identities of the cell bodies, which could be visualized on a Allen CCF 3D model brain (Fig. 1E).

### Higher visual areas, Auditory cortex, and Retrosplenial cortex provide strongest cortical inputs to V1

We found over fourteen brain regions of cortex and subcortex provided inputs to V1 (Fig. 2, Supplementary Fig. 3). In the thalamus, we found a dense cluster of labelled cells in dorsal lateral geniculate nucleus (dLGN) and lateral posterior nucleus (LP), which are visual areas known to provide inputs to V1 (Fig. 2B). In the cortex, we found labelled cells in higher visual areas (Fig. 2C) and other cortical areas such as retrosplenial cortex (RSP), anterior cingulate cortex (ACA) and medial entorhinal cortex (MEC) (Fig. 2D). In our samples with multiple injections in the same hemisphere, the spatial arrangement of the injections were echoed in LGN, LP and HVAs (Fig.2 B-C). We focused further analyses on non-thalamic regions whose inputs to V1 are less explored. To quantify the connection strength from each brain region, we calculated the percentage of the cells labelled by a given injection (excluding cells in V1, LGN and LP) occupied that brain region. We used percentage cells per injection to quantify the distribution of inputs to V1 to account for the variation in injection volume and efficacy of CTB transfection (raw cell counts across all areas are listed in Supplementary Table 3).

Higher visual areas had the most inputs to V1, followed by the retrosplenial, cingulate and other sensory cortices (Fig 3). The largest number of cells detected were in the HVAs (all HVAs: ~59%; median across all injections). Within the HVAs, we found the strongest inputs from the lateral visual area (L: 14.6%), followed by postrhinal cortex (POR: 8.6%) and posteriormedial (PM: 8.0%). In non-visual sensory areas we found highest cell counts in auditory areas (AUD: 6.6%), followed by somatosensory (SS: 2.7%) and motor (MO: 1.6%) areas. In non-sensory areas, we found cells in the retrosplenial cortex (RSP: 16.5%), temporal association area (TEa: 5.3%), anterior cingulate area (ACA: 3.2%), medial entorhinal area (MEC: 1.8%), ectorhinal area (ECT: 0.6%) and lateral entorhinal area (LEC: 0.5%). In addition, we found labelled cells in non-visual subcortical areas including the claustrum (CLA:2.3%) and the subicular complex (SUBcom:1.1%). We did not find differences in the distributions of cells between unilateral and bilateral injections (not shown), suggesting that bilateral injections did not result in labelling of callosal projections.

**Figure 3.**
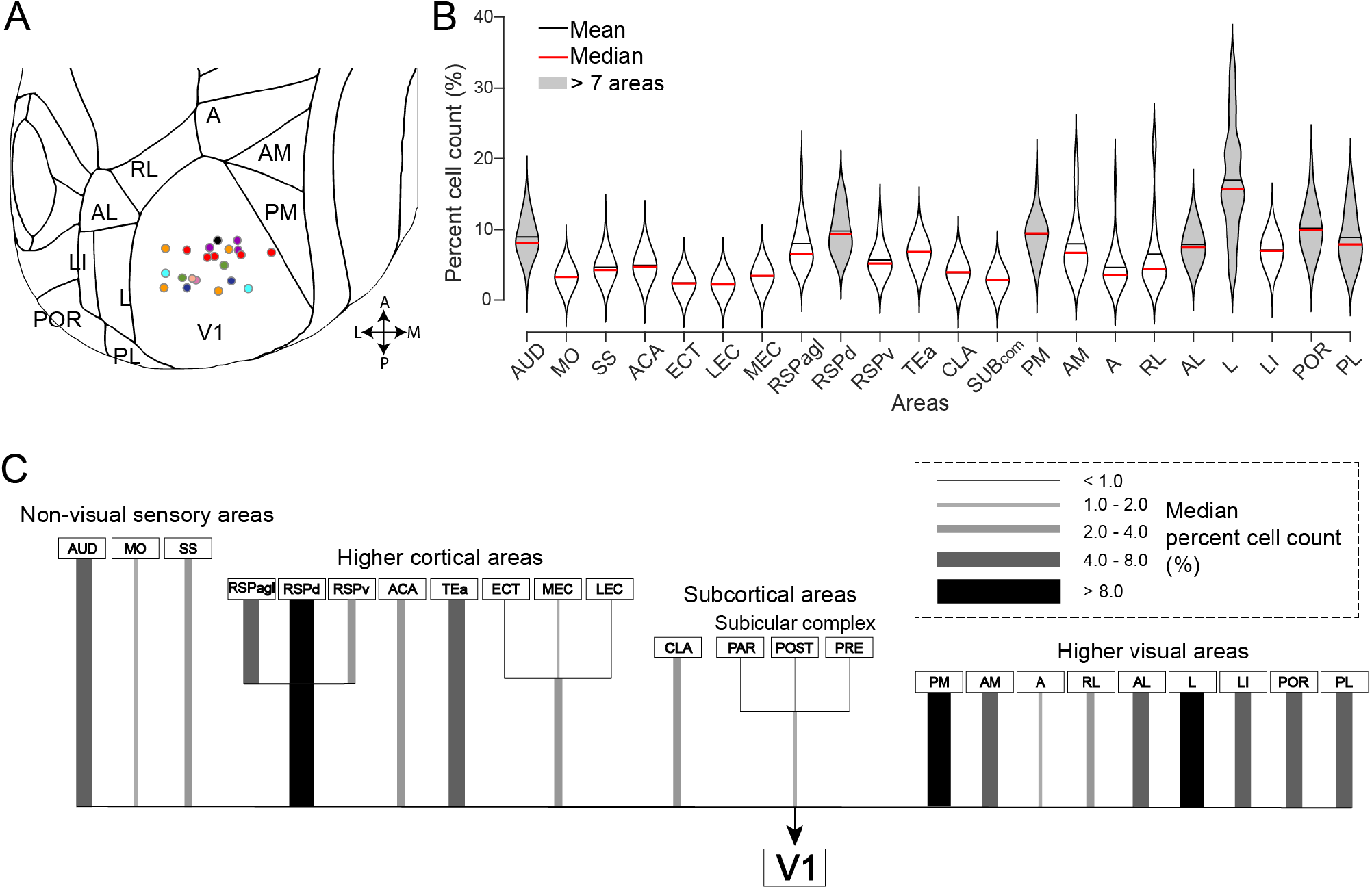
Distribution of cortical inputs to V1. **A)** Coordinates of all injection sites registered to the Allen Reference Atlas (21 injections; 9 animals). The colour of the dots represents which animal each injection belongs to. **B)** Distribution of percentage cell counts per area pooling all injections. (16 injections; 7 animals; mean: black line, median: red line). Gray fill indicates areas with significantly higher cell count than 7 other areas (kruskalwallis, p<0.05). For area name abbreviations see Supplementary Table 1. **C)** Data in B) illustrated as a tree diagram. Thickness and grayscale of lines relate to the median of percentage cell counts (%) as shown in legend.

The strength of input projections observed with our retrograde tracing methods was broadly consistent with results from anterograde tracing (Supplementary Fig. 4). We used projection data from the Allen Brain Connectivity Atlas to infer the strength of inputs into V1. We calculated the volume of fluorescent pixels in V1 based on anterograde tracing from all the source regions available. We found similar strength in the input to V1, with the HVAs being the dominant source of inputs. Other areas providing inputs to V1 included non-visual sensory areas, the retrosplenial cortex and the cingulate cortex (Supplementary Fig. 4). Note that quantifying the strength of innervation (axon terminals) with anterograde tracing can be noisy due to axons of passage, or thin axons below detection threshold. In addition, there was limited sampling as we only considered regions that had injections localised to within the region (see methods for criteria).

### Inputs from higher visual areas to V1 are topographically organized

As our injections into V1 spanned a range of stereotaxic coordinates (Supplementary Table 2), we asked if there was any organisation of the inputs into V1. In other words, do particular areas project preferentially to specific regions of V1? We first focused our analyses on HVAs, which had close to 60% of all labelled cells.

The distributions of cell counts from the different HVAs were quite variable between injections. Considering each injection site as an independent data point, we clustered the injection sites based on the distribution of cell counts in HVAs. We used k-means clustering and classified the data into two clusters. The clustering resulted in the injection sites being grouped into roughly medial and lateral clusters in anatomical space, even though no information about the injection site was used for clustering (Fig. 4A(i)). The separation into groups was visible when we viewed the first two principal components of distributions of HVA cell counts. (Fig. 4A(ii)). Splitting the data into additional clusters did not reveal further segregation. Therefore, injection sites broadly clustered into two groups, based on the distribution of inputs from different HVAs.

**Figure 4.**
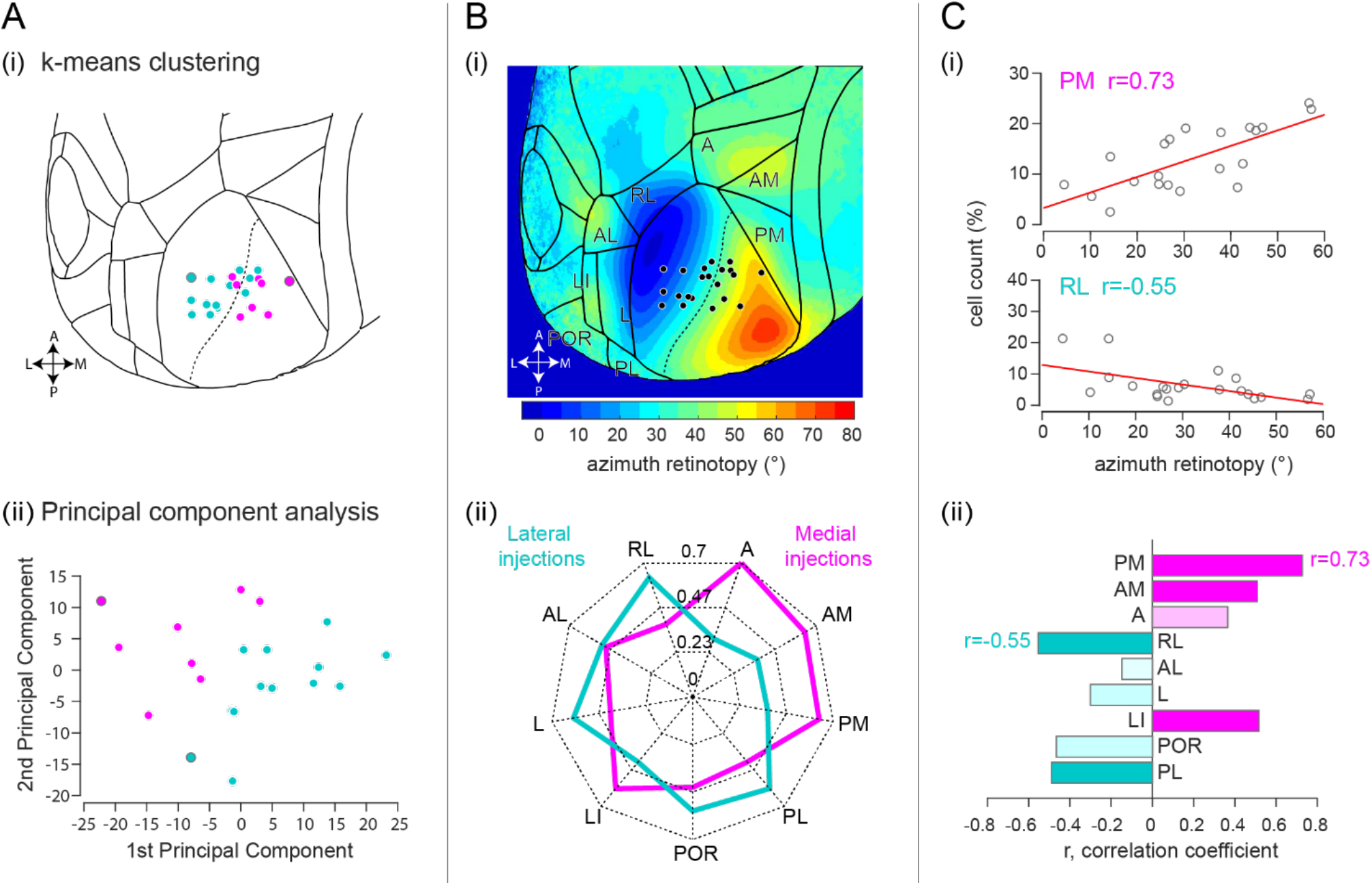
Input patterns from HVAs vary along the medio-lateral axis of V1. **A)** *Injection site clustering analysis:* (i) k-means clustering resulted in two groups corresponding to medial and lateral portions of V1. Points with gray outline denote data points used as seeds (21 injections; 9 animals). (ii) Applying this grouping in principal component space showed separation into two continuous groups. **B)** *Injection site grouping analysis:* (i) Injection coordinates plotted on mean azimuthal retinotopic map from Allen Institute. Contours are spaced 5° apart from −5° to 80° in azimuth. (ii) Grouping injections into medial and lateral groups show higher cell counts in lateral HVAs for lateral injections, and in medial HVAs for medial injections. **C)** *Per area correlation analysis:* (i) Correlation between azimuthal retinotopy of injection sites and cell count (shown for PM and RL). (ii) Positive correlations were observed for mostly medial areas (PM, AM, A, except LI), and negative correlations for lateral areas (RL, AL, L, POR and PL). Dark filled bars indicate significant correlations (p<0.05). Dotted lines in A-B correspond to 37.5° azimuth.

The grouping of injection sites based on distribution of cell counts in HVAs was consistent with a separation along retinotopy: into sites within the central visual field and the periphery visual field. To estimate the retinotopic location of each injection site, we overlaid the mean azimuthal map obtained from the Allen Institute dataset (Waters et al., 2019) and calculated the azimuthal position of each injection site (Fig. 4B, see methods). The separation into medial and lateral clusters was roughly along the 37.5° iso-azimuth line, which segregated the injection sites based on whether they were in the central or peripheral visual fields. Therefore, we grouped the injections based on being below (medial) or above (lateral) 37.5°. Based on this grouping, we found that medial injections had preferentially higher cell counts in medial HVAs (A, AM, PM), and lateral injections had higher cell counts in lateral HVAs (L, RL, PL, POR) (Fig. 4D).

The percentage of cells detected in HVAs were correlated to the azimuthal retinotopic location of the injection site. We also assessed the relationship between the retinotopic location of injection sites and distribution of cell counts in each HVA by plotting the estimated azimuthal coordinates of all injection sites against their corresponding cell counts (Fig. 4C, Supplementary Fig. 5). We observed significant correlations between the cell counts and azimuth across many HVAs, with predominantly positive correlations for HVAs that were medial to V1 (PM, AM A), and negative correlations for most lateral areas (RL, POR, PL, with the exception of LI). Therefore, both correlation and clustering results indicate that there is a difference in input pattern along the mediolateral axis of V1, which suggests that there is a topographic organization of feedback inputs from HVAs to V1.

The topographic organisation of inputs into V1 was significant only for the HVAs and not for other cortical regions of the brain (Fig. 5). Given the topographic organisation in HVA inputs to V1, we next analysed inputs to V1 from all brain regions using the split of injection sites into medial and lateral groups (Fig. 5A). We found minor variations in input pattern across nonvisual brain areas (Fig. 5B). Only HVAs, specifically RL, AL, L, POR and PL, showed a significant difference between medial and lateral injection groups (Fig. 5B). This suggests that the mediolateral bias in inputs to V1 might not extend beyond visual areas.

**Figure 5.**
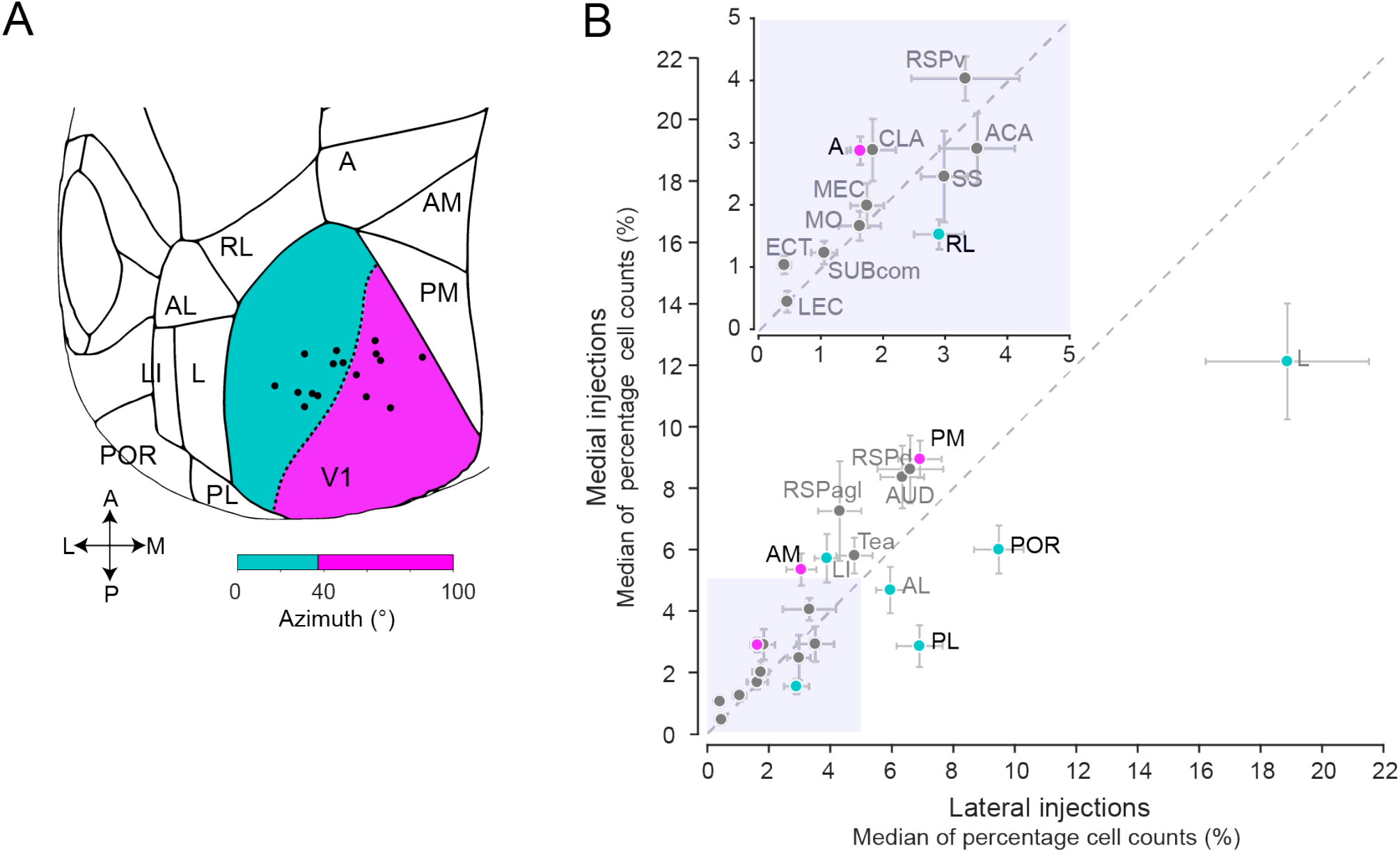
Topographic organization of inputs to Medial vs Lateral V1. **A)** Grouping of injections into medial and lateral groups based on retinotopic coordinates (16 injections; 7 animals). Lateral group indicated in cyan (azimuth 0° to 37.5°), medial group indicated in magenta (azimuth 37.5° to 100°). **B)** Median of normalized cell counts (%), medial and lateral injection groups plotted against each other. Error bars indicate S.E.M.. Gray dotted line indicates the unity line. Lateral HVAs in cyan, medial HVAs in magenta. Names of areas with significant difference between medial and lateral injection groups are in black (Wilcoxon rank sum test, p<0.05). Shaded region (0 to 5) enlarged in inset.

## Discussion

In this study, we investigated cortex-wide inputs to mouse V1 using the retrograde tracer CTB. We first developed a simple and modular software pipeline for the quantification of labelled cell bodies in Allen CCF, which identifies the brain region of each labelled cell. Confirming previous reports, and also in agreement with anterograde data from the Allen Brain Institute, we observed cortex-wide inputs to V1, including higher visual areas, retrosplenial cortex, other sensory cortices, cingulate cortex and rhinal areas. We next quantified inputs from across the brain. The most inputs, approximately 60% of detected cells, were from higher visual areas (HVAs). Within HVA cells, we discovered a topographic organization in their projection to medial vs lateral V1. Medial HVAs (A, AM, PM) preferentially project more to medial V1, whereas lateral HVAs (RL, AL, L, POR, PL, but not LI) preferentially project more to lateral V1.

### Software pipeline for quantifying retrogradely labelled cells

We developed a software pipeline to quantify and annotate cells based on the now standard mouse brain map, Allen Reference Atlas (Wang et al., 2020). While other software has recently been developed for this purpose (Fürth et al., 2018; Song et al., 2020; Tyson et al., 2020), our strategy is simple and modular, with little learning curve. It is semi-automated, and requires human curation for cell detection and alignment to the Allen CCF. Our software takes advantage of sharp-Track (Shamash et al., 2018) and has been developed as an extension of that framework, so that the same software package can be used for tracking electrode positions and detecting cell-bodies.

The definition of brain areas in our study was based on the Allen Reference Atlas, coregistered to our brain images in Allen CCF. The accuracy of area identification using this method was previously shown to be better than visual inspection (Shamash et al., 2018). As a sanity check, we verified whether known architectonic signatures in our DAPI images matched our area annotations, such as rapid thinning of layer 4 as the transition between retrosplenial cortex and HVAs and found them to be consistent with each other.

### Diverse areas projecting to the primary visual cortex

We found fourteen unique non-visual brain regions that project to V1. These results suggest that top-down inputs associated with V1 activity might be routed directly from these areas rather than indirectly through HVAs. For example, we find inputs from the auditory areas (AUD), which might be involved in generating the multi-sensory integration observed in V1 (Iurilli et al., 2012). Inputs from the anterior cingulate cortex have been attributed to modulate locomotor signals (Fiser et al., 2016; Leinweber et al., 2017), and those from the claustrum could be involved in change detection and modulating the salience of visual cues (Brown et al., 2017; Atlan et al., 2018). Spatial signals modulating V1 activity (Pakan et al., 2018; Saleem et al., 2018; Fournier et al., 2019) could be routed through areas detected that are also known to have spatial signals, including the retrosplenial cortex (Witter et al., 2017; Mao et al., 2018; Nitzan et al., 2020), or the entorhinal and subicular areas (Witter et al., 2017).

Using this pipeline, we detected high cell counts in HVAs, especially L, PM, AL and POR, dorsal retrosplenial cortex (RSPd) and auditory cortex (AUD). This ranking of inputs per area is consistent with previous studies using retrograde tracing, albeit minor differences that may have been caused by use of different reference maps (Zingg et al., 2014; Leinweber et al., 2017). We also saw in our double or triple injection experiments that double or triple labelled cells were rarely found (data not shown). This suggests that single cortical cells do not provide widespread input across V1 spanning our multiple injection sites.

### Topographic organization of feedback inputs from HVAs

A defining feature of mouse HVAs is that each area has a retinotopic representation of the visual space, or retinotopic map. The retinotopic maps in HVAs have been shown to represent visual space in a biased manner, with close by regions representing similar regions of visual space (Zhuang et al., 2017). This mesoscale organization is likely a result of topographically organized feedforward projections from V1 to HVAs.

Here, we observed that inputs from HVAs to V1 are topographically organized. This is largely consistent with some studies that investigated specific projections from HVAs to V1. For example, feedback projections from LM have matched receptive fields to V1 cells in the vicinity of its projections (Marques et al., 2018), and AL and PM feedback projections were shown to differentially affect V1 neurons when silenced (Huh et al., 2018).

In ferrets and macaques, it is known that feedback from HVAs show retinotopic convergence, in which HVA feedback neurons represent a larger retinotopic area compared to the V1 region it provides feedback to (Angelucci et al., 2002; Cantone et al., 2005). These feedback projections are therefore likely to contribute to surround suppression in V1, which was shown to be the case using optogenetics in marmosets (Nurminen et al., 2018) and mice (Vangeneugden et al., 2019). From our data it was not possible to understand how much of the topography we observed could be explained by the retinotopy of V1 recipient neurons and HVA feedback neurons. This was because we did not empirically measure the retinotopic location of the HVA somata and terminals, but also because our injection volume in V1 was relatively large and spanned a large extent of retinotopic space. Thus, it would be of interest for future studies to trace single cell inputs retrogradely to understand the precise retinotopy of HVA inputs to V1.

### Functional implications of topographic organization of feedback inputs

Growing evidence points to functional diversity within the V1 population. Binocular disparity tuning, colour tuning, and coherent motion processing have recently been shown to differ along the retinotopic elevation axis of V1 (Aihara et al., 2017; La Chioma et al., 2019; Sit and Goard, 2019). Moreover, in a mouse performing a task in virtual reality, task related variables were observed to be represented continuously across visual areas (Minderer et al., 2019). These studies argue for a topological organization of mouse visual areas at the functional level.

The topographic organization of inputs we observed between medial and lateral V1 might mean that V1 cells representing peripheral vs central visual fields are differentially modulated by HVAs. Ethologically, animals encounter different types of visual information in different portions of the visual field. In the case of a navigating mouse, its peripheral vision might be more frequently used for detecting optic flow, whereas the central vision might be used for landmarks to orient to. This has led to the proposal for a central and peripheral stream of processing for mouse vision (Saleem, 2020), and the current finding for HVA to V1 feedback projections is consistent with this notion. This view is further supported by recent findings that different HVA projections convey different information to V1 cells (Huh et al., 2018), and that these feedback projections follow a ‘like-to-like’ rule in their reciprocal connection to V1 cells (Marques et al., 2018; Kim et al., 2020).

However, such functional proposals need to be further assessed in the future, taking into account recurrent connections within V1, and connection between HVAs. With more knowledge about the cell types of these feedback projections and what information they convey during behaviour, we will have a better understanding of the functional implication of these mesoscale organizations.

## Acknowledgements

We thank Thomas Wheatcroft for sharing code for pre-processing brain images. We also thank Stefano Zucca, Philip Shamash, Andrew MacAskill, Rawan AlSubaie and Candela Sánchez-Bellot for advice and useful discussions. We appreciate Jack Water’s guidance on alignment of Allen Institute’s retinotopic maps to our dataset. Finally, we thank all members of the lab and Samuel Solomon for comments and discussions. This work was supported by the Wellcome Trust (200501), Biotechnology and Biological Sciences Research Council (R004765), HFSP (RGY0076/2018), and Royal Society (RGS\R1\191449) grants.

## Author Contributions

This study was conceptualized by MMM and ABS; the methodology was developed by MMM; the data was collected, curated and analysed by MMM and EU; the visualization and software was by MMM, EU and ABS; the project was supervised by MMM and ABS; funding was acquired by ABS; and the article was written by all authors.

## Data and Software Availability statement

The processed data will soon be available to download at *FigShare*, and the code at *Github*. Original images of brain slices are available upon reasonable request.

## Materials & Methods

### Surgery and injection

All procedures were conducted in accordance with the UK Animals Scientific Procedures Act (1986). Experiments were performed at University College London under personal and project licenses released by the Home Office following appropriate ethics review.

We used nine adult male and female wild-type mice (C57BL/6J, aged 13–30 weeks) for this study. Mice were anaesthetised with 2% isoflurane delivered with oxygen (0.5 l/min), and placed on a heating pad to maintain body temperature. An incision was made to the scalp along the midline to expose the injection area, and injection coordinates for V1 (listed in Supplementary Table 2) were marked using a stereotaxic software (Robot Stereotaxic, Neurostar). Small craniotomies were made at these sites and cholera toxin subunit B (CTB) conjugated with Alexa Fluor (CTB-488, −555, −647, Thermo Fisher) was injected using glass micropipettes. CTB was injected in 2 to 3 sites in V1 bilaterally (7 mice) or unilaterally (2 mice) in V1, at depths of 250μm or 750μm below the pia (unilateral: M19118, M19119, bilateral double: M19114, M19115, M19121, M19122, M19123 bilateral triple: M19116, M19117, see Supplementary Table 2 and Supplementary Fig. 1 for more details). Multiple injections within a hemisphere were made with unique fluorophores (CTB-488, −555, and −647). At each site, we infused 200-300nl at a flow rate of 20- 40nl/min. The micropipette was left at the injection site for 5 minutes after the infusion completed, before being withdrawn slowly. We covered the craniotomies with KwikCast (World Precision Instruments), sutured the scalp, and allowed the animals to recover for 3 days while orally administering analgesics.

### Histology and imaging

10-14 days post-injection, mice were anesthetized with 5% isoflurane and injected with pentobarbital intraperitoneally. Mice were transcardially perfused with 0.9% NaCl in 0.1M phosphate buffer (PB), followed by 4% paraformaldehyde (PFA). The brain was extracted and placed in 4% PFA overnight at 4°C, and subsequently cryoprotected with 30% sucrose solution. The olfactory bulb and cerebellum were cut away, and brains were frozen in O.C.T. Compound (Sakura FineTek). Coronal sections of 50μm thickness were sliced on a cryostat (Leica, CM1850 UV), on average between AP-coordinates of bregma 0.64±0.30 (S.E.M.) to −4.55±0.08 (S.E.M.). Slices were mounted using Vectorshield with DAPI (Vector Labs) or ProLong Diamond Antifade Mountant with DAPI (Invitrogen) and slices 150μm apart were imaged with a standard fluorescence microscope (Leica DMi8 or ZEISS Axio Scan.Z1) using a 10x objective and standard filter sets. The contrasts of individual channels were adjusted in figures to optimise visualisation.

### Detection and quantification of labelled cells

Our analysis pipeline consisted of a cell detection procedure followed by alignment to the Allen CCF using SHARP-Track (Shamash et al., 2018). Our cell detection algorithm steps (implemented using the image processing toolbox in MATLAB, R2017; function names in italic) were: **1)** background subtraction using morphological top-hat filter ***imtophat*** (structural element:’disk’, size: 4), **2)** binarization by ***imbinarize*** using a percentile threshold (typically 99.5 to 99.8 percentile of the fluorescence intensity histogram) 3) selecting objects larger than 25 pixels in the binary image using ***bwareaopen*, 3)** erode and dilate edges of objects using ***imerode/imdilate*** (structural element: ‘disk’; size: 1) **4)** fill holes in the objects through ***imfill*** followed by **5) *watershed*** to isolate connected objects, and finally **6)** extract centroids of objects using ***regionprops*** (property: area > 20). Using the centroids from the final step, we generated a centroid mask (an image that has a value of 1 at the centroid locations and 0 elsewhere) of the same size as the original image and used this and DAPI channels as input to SHARP-Track. As SHARP-Track was originally developed to analyse electrode tracks, we adapted some functions for the purpose of quantifying cells in different brain regions. The GUI in SHARP-Track allows for the user to scroll through slices of the Allen Reference Atlas (ARA) 3D brain model, and visually identify the slice that best corresponds to the imaged brain slice (DAPI image). The selected ARA slice (reference slice) can be micro-adjusted in the AP-, ML- and DV-axes, to account for any asymmetry introduced during histological procedures. Subsequently, transformation points were selected by clicking on corresponding anatomical landmarks between the DAPI image and reference slice. These points are used to morph the DAPI image to fit the reference slice, using local transformations (Shamash et al., 2018). We placed more transform points around our areas of interest (e.g. where we found cells) such that the fit would be more accurate around the areas of interest. We then used the same transformation on the centroid mask image to map cell body centroids to the ARA. This generated a table of all detected cells with coordinates and area identities in Allen CCF (based on the Allen Reference Atlas). We analysed each of the imaged slices through this pipeline and calculated the cell counts from each injection site. Absolute cell counts were normalized by the total number of cells per injection (excluding cells in V1, LGN and LP). Cell counts from individual injection sites were considered independent samples for subsequent analyses. For estimating the centroid of injection sites in Allen CCF, we first used image processing methods almost identical to cell detection described above. For each injection, we identified the injection site in the slice with the strongest fluorescence signal (Supplementary Fig. 1), then after segmenting the injection site as a single object, centroids were extracted using regionprops (property: area > 5000). Subsequent steps to align the centroid to ARA was identical to the method described above.

### Retinotopic location of injection sites

To estimate the retinotopic position of our injection sites post hoc, we followed a procedure similar to Minderer et al. 2019, and used intrinsic imaging data published in (Waters et al., 2019). This dataset contained mean azimuth and elevation maps in Allen CCF from 60 mice, generated using horizontal and vertical sweeping bars (Kalatsky and Stryker, 2003). Using the injection site coordinates in Allen CCF, obtained using the software pipeline above, we directly read out values from the mean azimuthal map and used these as estimated retinotopic positions of our injection sites. While we did not experimentally confirm the individual retinotopic location of each injection site, previous work has shown that the difference in the retinotopic maps across animals is comparable to measurement error across trials within individual animals, due to the underlying variability in the estimation of the receptive field using wide-field intrinsic imaging (Waters et al., 2019).

### Analysis of anterograde tracing data

We used anterograde tracing data from the publicly available connectivity repository of the Allen Brain Institute (https://connectivity.brain-map.org/projection). Specifically, we used the ‘target search’ functionality in the GUI provided at the website. We selected specific brain regions (shown in Supplementary Fig. 4) as ‘source’, and V1 (‘VISp’) as the ‘target’ and analysed all resulting injections from all mouse strains. The data contained measured values such as the volume of fluorescent pixels (mm^3^) in the target region, and injection specificity. We adopted a criterion to filter these injections: selecting for further analyses only those with injection volumes over 0.1 mm^3^, and injection specificity over 70%. We then normalized the fluorescent volume of the target region by the injection volume to account for the variability in injection volumes. This allowed us to assess the amount of specific projections from ‘source’ areas to ‘target’ area V1 using anterograde tracing data.

### Other Analyses

To cluster injections based on the distribution of inputs connections, we used k-means clustering. For this clustering, we used two seeds, one in medial V1 and another in lateral V1. This was necessary because the seeds needed to be spaced apart sufficiently to produce meaningful groups. For correlation analysis, we used the Pearson’s correlation coefficient r.

Paired t-tests and Kruskal-Wallis tests were used to test for statistical significance. For all comparisons, we used a significance level of p < 0.05.

All analyses were performed in MATLAB.

## Supplementary Tables

**Supplementary Table 1:** Area name abbreviations used in the study. Most follow Allen CCF (v3) nomenclature (Wang et al., 2020). Exceptions are indicated in italic, and corresponding Allen CCF (v3) nomenclature shown in brackets where existing.

|  |  |
| --- | --- |
| *V1* (VISp) | Primary visual area |
| AUD | Auditory areas |
| MO | Somatomotor areas |
| SS | Somatosensory areas |
| RSP | Retrosplenial area |
| RSPagl | Retrosplenial area, lateral agranular part |
| RSPd | Retrosplenial area, dorsal part |
| RSPv | Retrosplenial area, ventral part |
| ACA | Anterior cingulate area |
| TEa | Temporal association areas |
| ECT | Ectorhinal area |
| *MEC* (ENTm) | Entorhinal area, medial part, dorsal zone |
| *LEC* (ENTl) | Entorhinal area, lateral part |
| CLA | Clastrum |
| SUBcom | Subicular complex |
| PAR | Parasubiculum |
| POST | Postsubiculum |
| PRE | Presubiculum |
| *PM* (VISpm) | Posteromedial visual area |
| *AM* (VISal) | Anteromedial visual area |
| *A* (VISa) | Anterior area |
| *RL* (VISrl) | Rostrolateral visual area |
| *AL* (VISal) | Anterolateral visual area |
| *L* (VISl) | Lateral visual area |
| *LI* (VISli) | Laterointermediate area |
| *POR* (VISpor) | Postrhinal area |
| *PL* (VISpl) | Posterolateral visual area |

**Supplementary Table 2:**
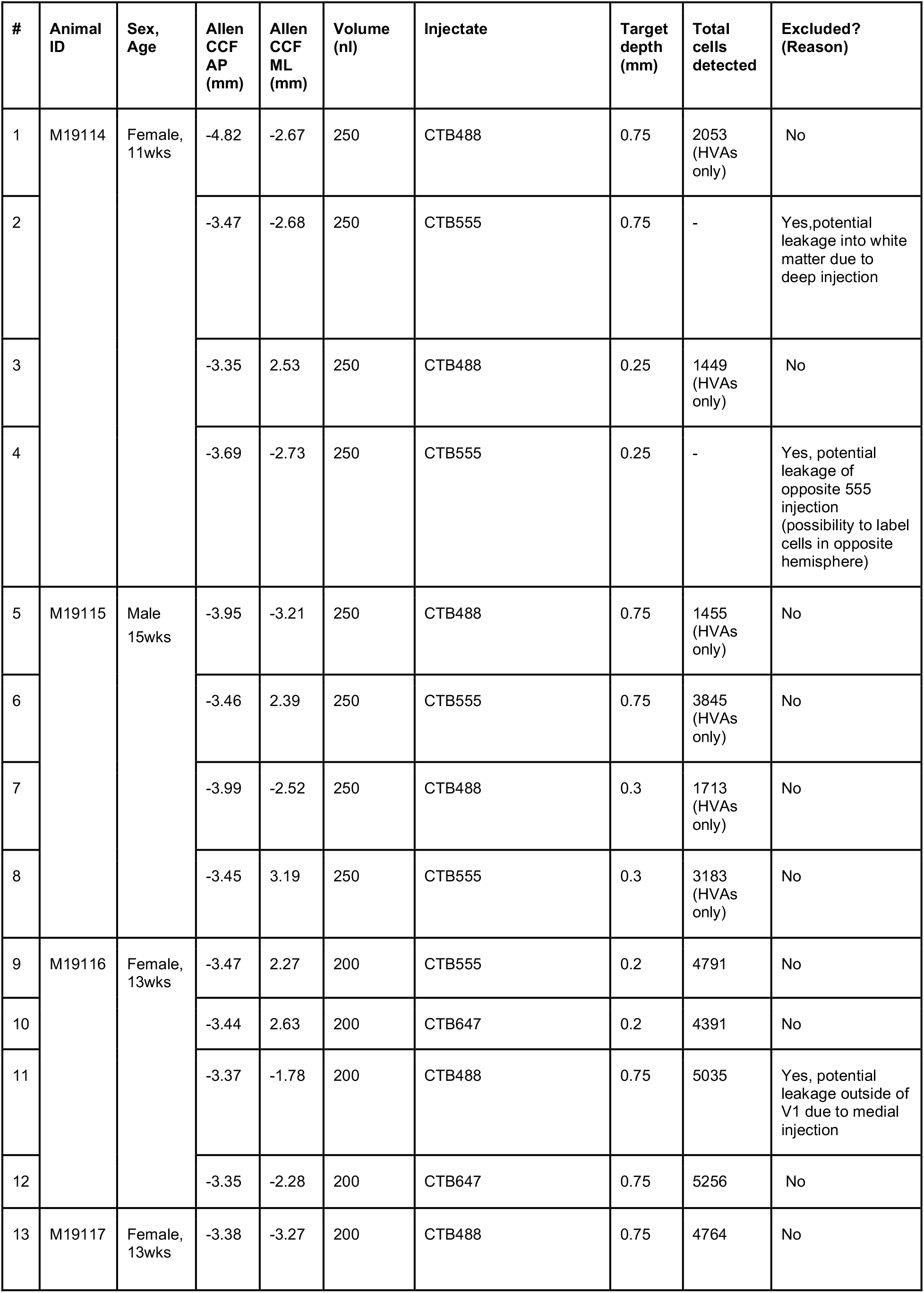
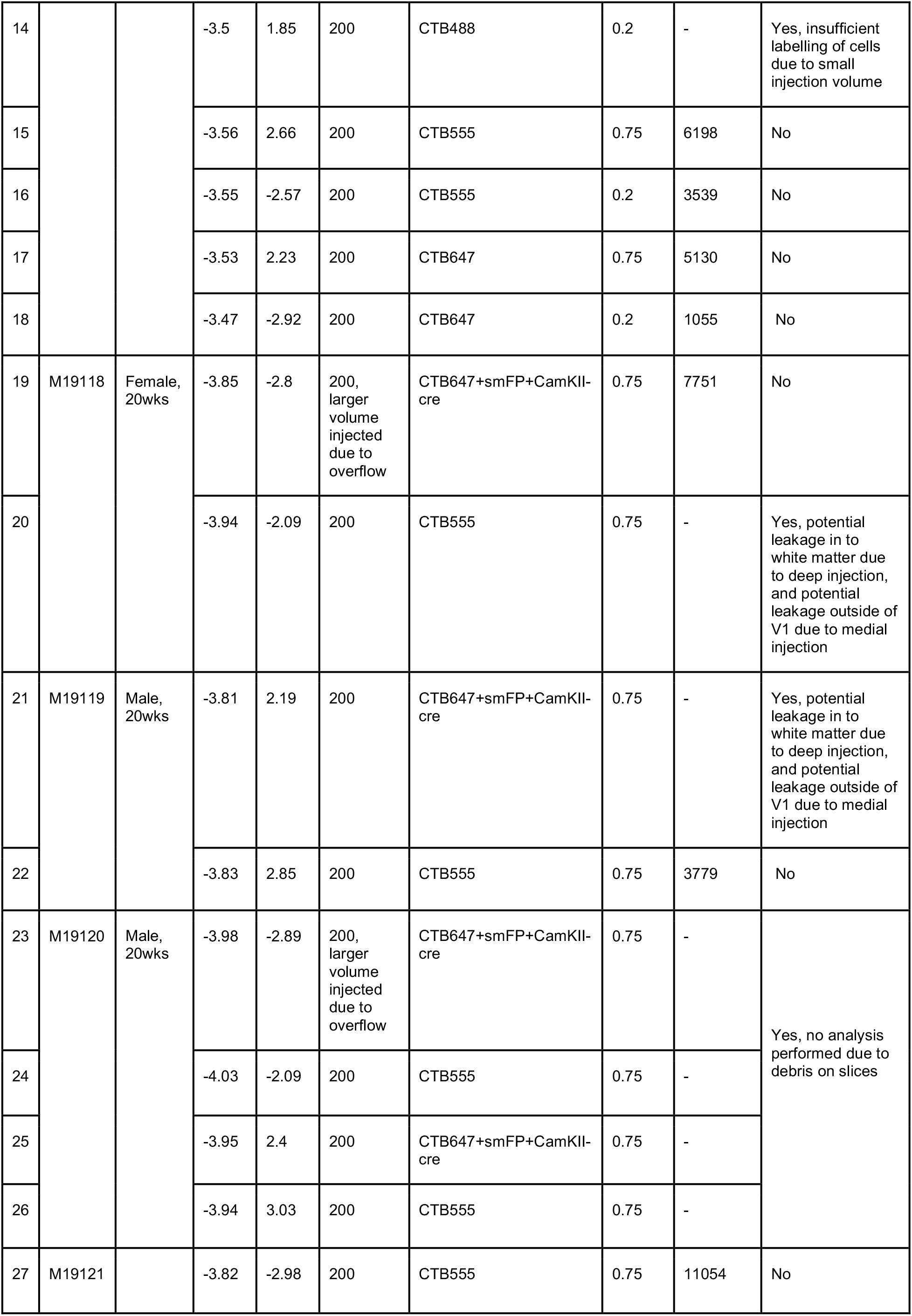
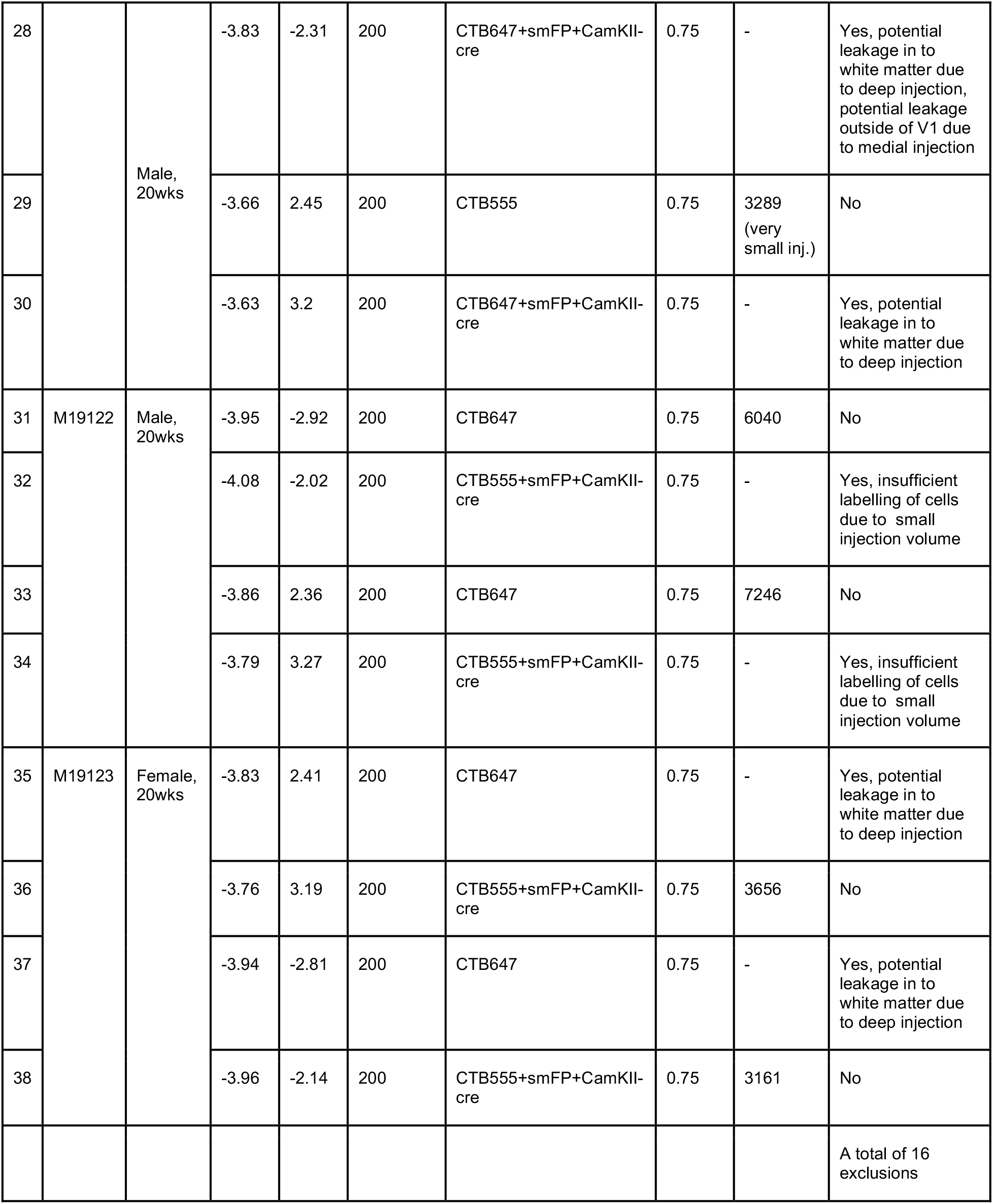
List of all injections in the study, including injection coordinates, injectate, and total number of cells detected

**Supplementary Table 3:** Number of cells detected across all brain regions.

| Area | Relative cell count |  |  | Absolute cell count |  |
| --- | --- | --- | --- | --- | --- |
|  | Mean | SEM | Median | Total cells counted | from 'n' injections |
| AUD | 7.465 | 0.679 | 6.615 | 5895 | 16 |
| MO | 1.606 | 0.224 | 1.645 | 1464 | 16 |
| SS | 3.063 | 0.402 | 2.652 | 2543 | 16 |
| ACA | 3.332 | 0.433 | 3.172 | 2777 | 16 |
| ECT | 0.669 | 0.098 | 0.601 | 537 | 16 |
| LEC | 0.545 | 0.092 | 0.457 | 476 | 16 |
| MEC | 1.750 | 0.227 | 1.804 | 1558 | 16 |
| RSPagl | 6.484 | 0.954 | 4.975 | 5144 | 16 |
| RSPd | 8.419 | 0.794 | 7.906 | 6714 | 16 |
| RSPv | 4.107 | 0.534 | 3.588 | 3255 | 16 |
| Tea | 5.279 | 0.437 | 5.284 | 4220 | 16 |
| CLA | 2.270 | 0.331 | 2.320 | 2063 | 16 |
| SUBcom | 1.197 | 0.139 | 1.171 | 990 | 16 |
| PM | 7.837 | 0.648 | 7.988 | 6991 | 21 |
| AM | 6.467 | 1.066 | 5.163 | 5317 | 21 |
| A | 3.039 | 0.712 | 1.908 | 2531 | 21 |
| RL | 4.965 | 1.245 | 2.780 | 3348 | 21 |
| AL | 6.358 | 0.521 | 5.940 | 5698 | 21 |
| L | 15.940 | 1.748 | 14.581 | 14095 | 21 |
| LI | 5.575 | 0.440 | 5.458 | 4814 | 21 |
| POR | 8.830 | 0.787 | 8.573 | 7470 | 21 |
| PL | 7.401 | 0.865 | 6.389 | 5898 | 21 |

## Supplementary Figures

**Supplementary Figure 1:**
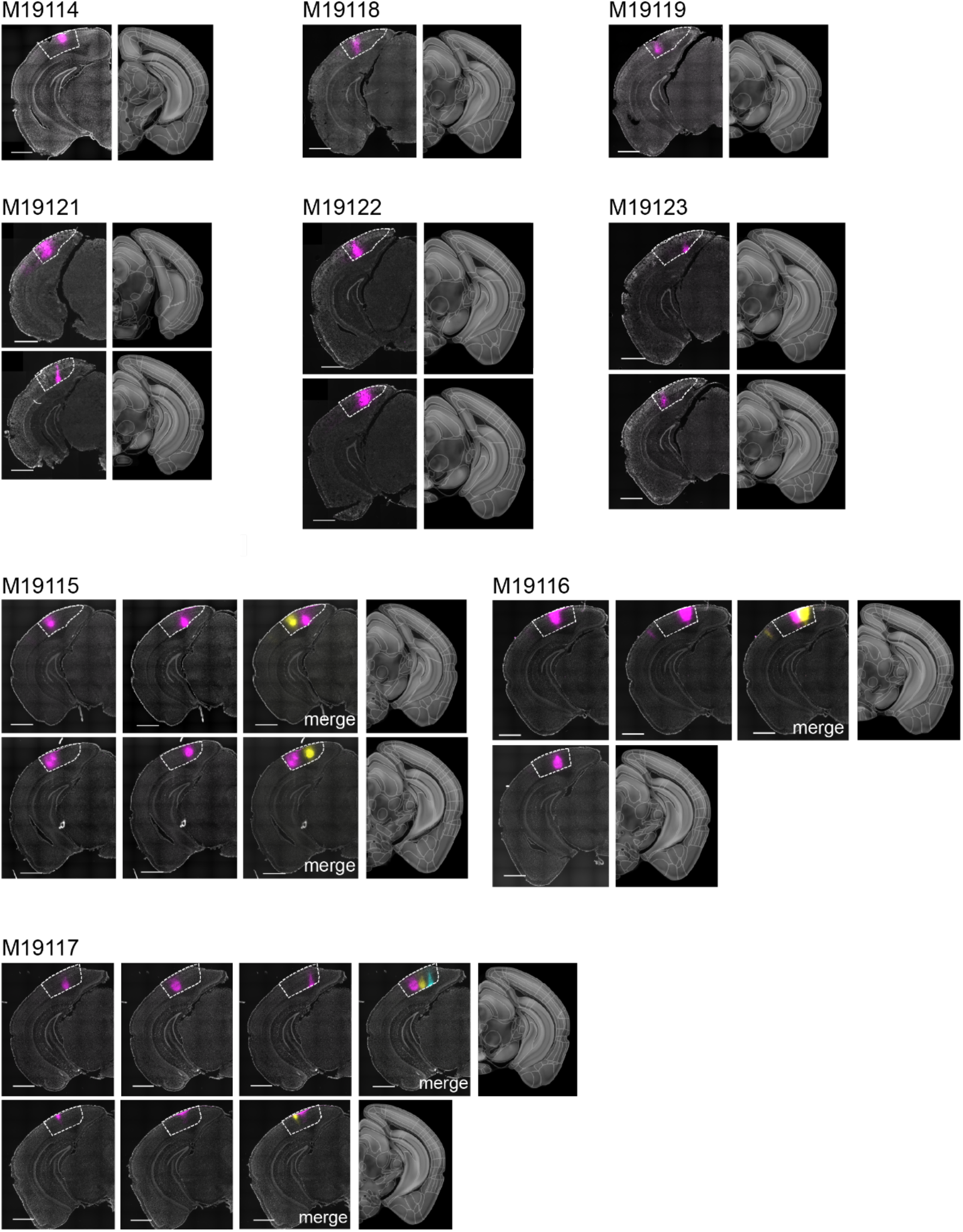
CTB injection sites used in the study (9 animals, 21 injections). Fluorescent signal from each injection is shown in magenta. In ‘merge’ images for multiple injections within the same hemisphere, additional injections are shown in yellow or cyan. For each brain image (left hemisphere image), the Allen Reference Atlas (ARA) image used for alignment is shown on the same row (right hemisphere image). The brain images are pre-morphing, before alignment to the reference image. Border of V1 is indicated in white dotted lines according to this ARA image. Scale bars = 1mm. Further details of injections in methods and Supplementary Table 1.

**Supplementary Figure 2:**
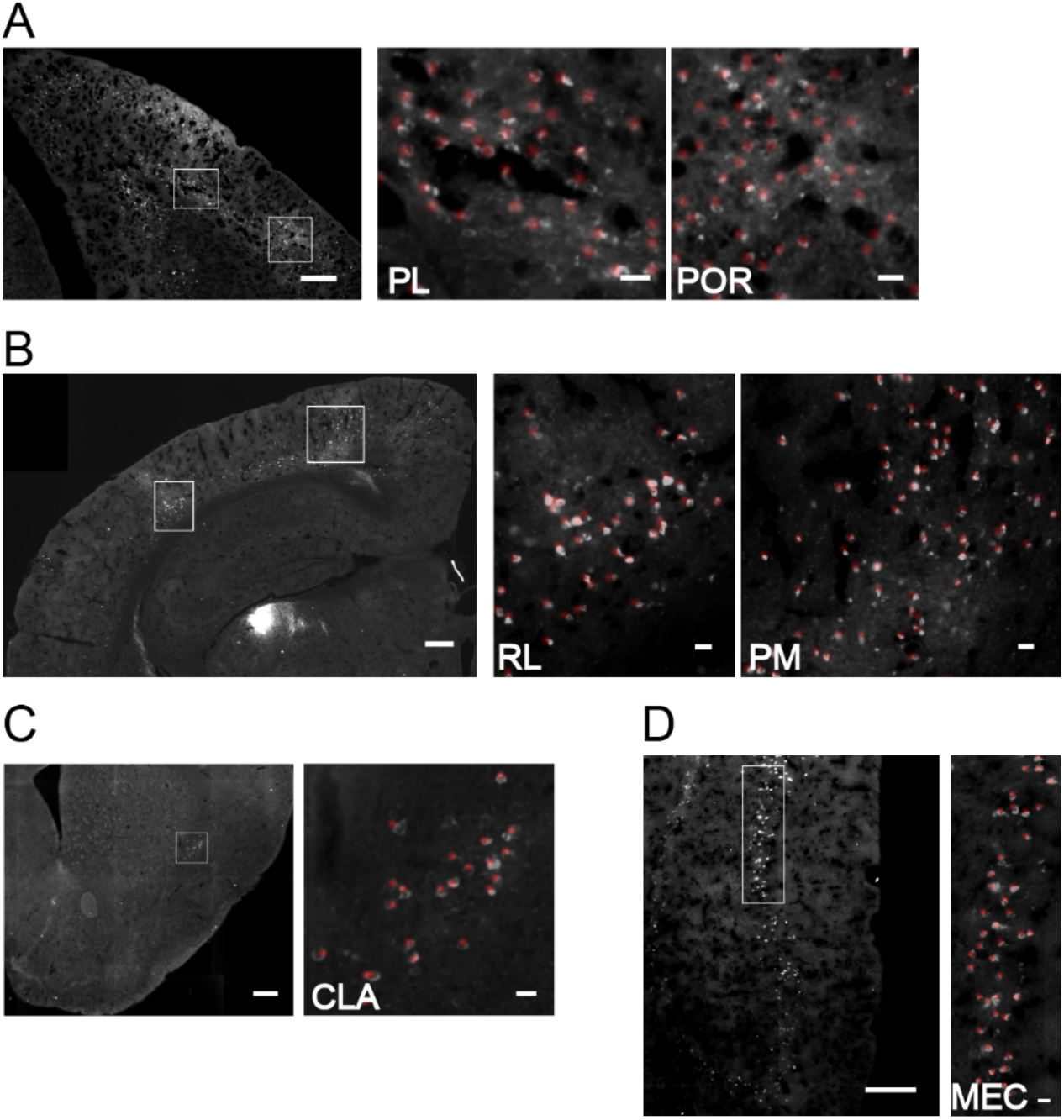
**A-D)** Cell detection through our software pipeline. Red regions indicate detected cells. Right panels correspond to white boxed regions on the left panel. Scale bars: left panel = 100μm, right panel = 10μm.

**Supplementary Figure 3:**
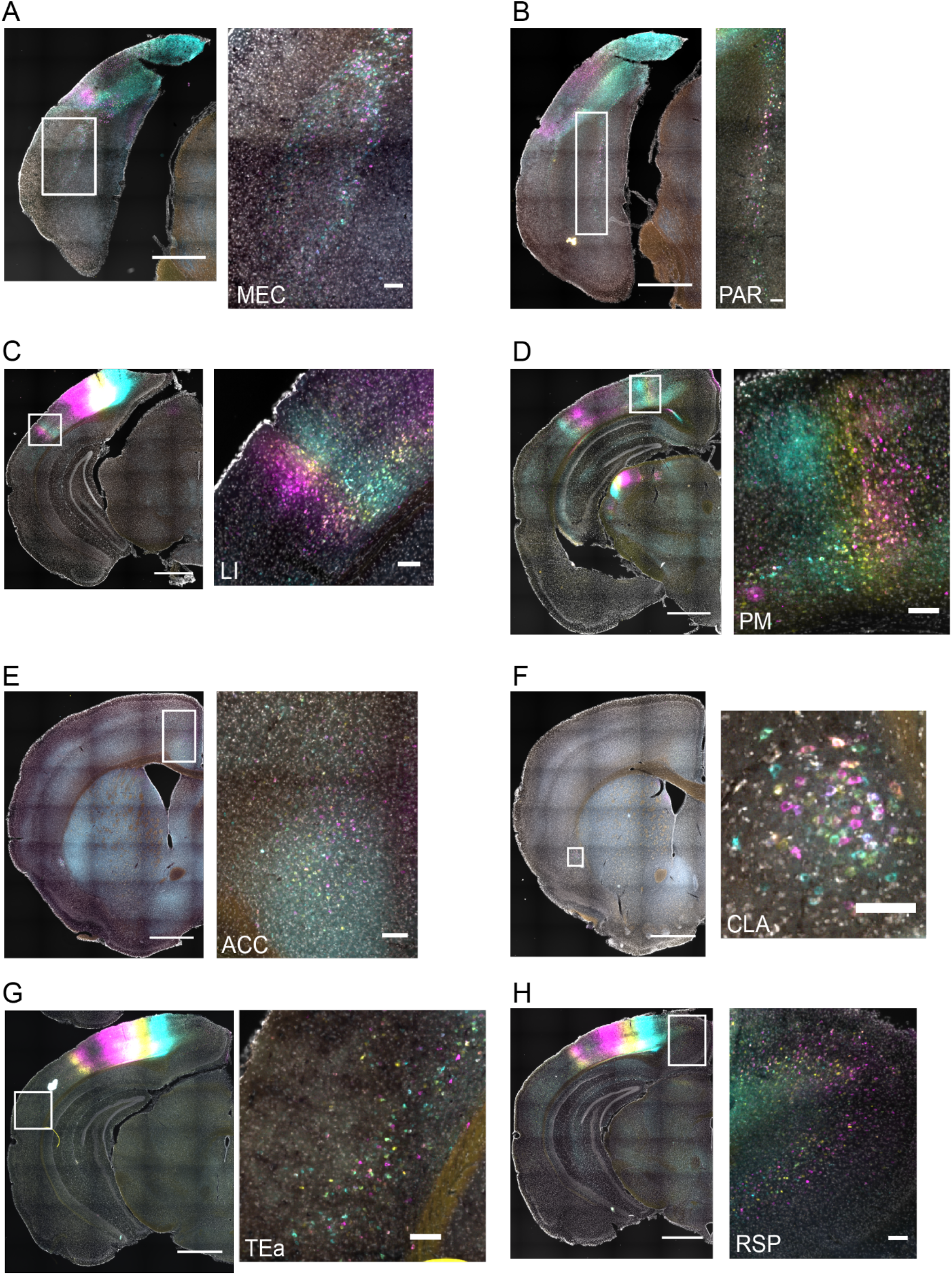
Retrogradely labelled cells detected in various brain regions. **A-H)** Example areas showing retrogradely labelled cells. Right panels correspond to white boxed regions on the left panel. Scale bars: left panel = 1 mm, right panel = 100 μm.

**Supplementary Figure 4:**
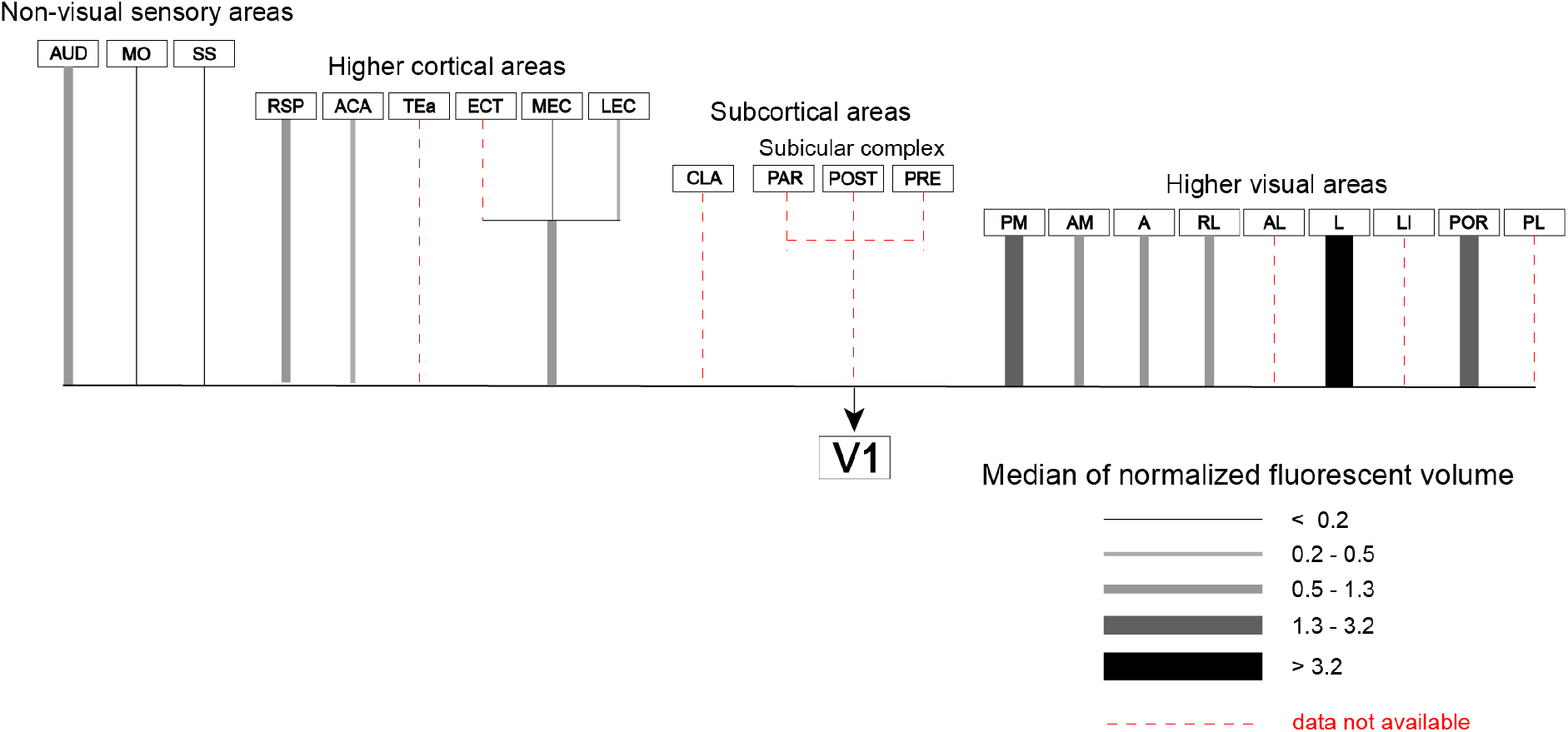
Estimated projection to V1 from brain wide anterograde injection data from the Allen Brain Institute. Injections to these areas were curated for specificity and projection to V1 from each injection was quantified (volume of fluorescent pixels in V1 normalized by injection volume). See methods for further details.

**Supplementary Figure 5.**
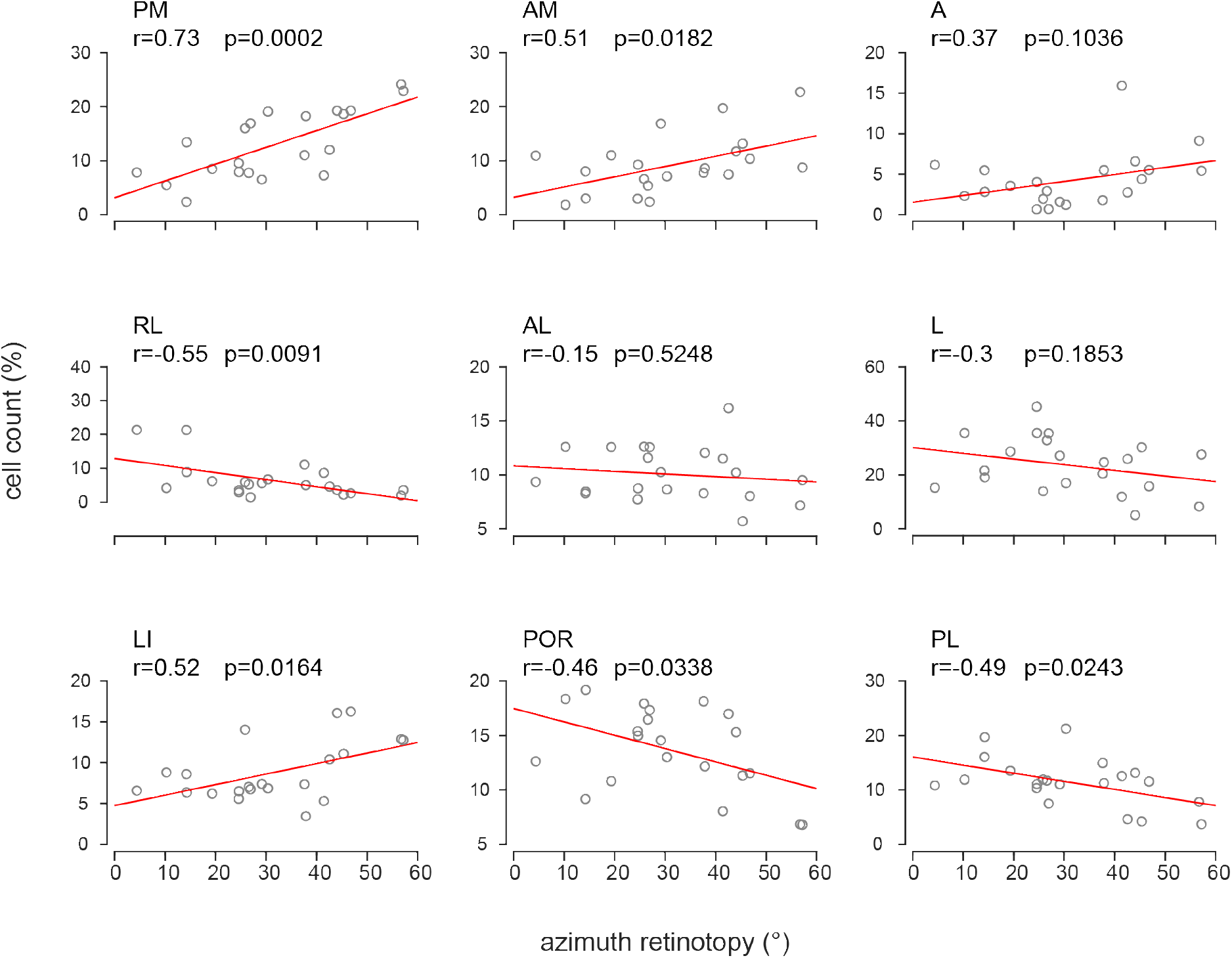
(related to Figure 4C): Correlation between estimated azimuth retinotopy of injection sites and detected cell counts in each HVA.

## References

Aihara S, Yoshida T, Hashimoto T, Ohki K (2017) Color representation is retinotopically biased but locally intermingled in mouse V1. Front Neural Circuits 11:22.

Andermann ML, Kerlin AM, Roumis DK, Glickfeld LL, Reid RC (2011) Functional specialization of mouse higher visual cortical areas. Neuron 72:1025–1039.

Angelucci A, Levitt JB, Walton EJS, Hupe J-M, Bullier J, Lund JS (2002) Circuits for local and global signal integration in primary visual cortex. J Neurosci 22:8633–8646.

Atlan G, Terem A, Peretz-Rivlin N, Sehrawat K, Gonzales BJ, Pozner G, Tasaka G-I, Goll Y, Refaeli R, Zviran O, Lim BK, Groysman M, Goshen I, Mizrahi A, Nelken I, Citri A (2018) The claustrum supports resilience to distraction. Curr Biol 28:2752–2762.e7.

Brown SP, Mathur BN, Olsen SR, Luppi P-H, Bickford ME, Citri A (2017) New Breakthroughs in Understanding the Role of Functional Interactions between the Neocortex and the Claustrum. J Neurosci 37:10877–10881.

Cantone G, Xiao J, McFarlane N, Levitt JB (2005) Feedback connections to ferret striate cortex: direct evidence for visuotopic convergence of feedback inputs. J Comp Neurol 487:312–331.

Fiser A, Mahringer D, Oyibo HK, Petersen AV, Leinweber M, Keller GB (2016) Experience-dependent spatial expectations in mouse visual cortex. Nat Neurosci 19:1658–1664.

Fournier J, Saleem AB, Diamanti EM, Wells MJ, Harris KD, Carandini M (2019) Modulation of visual cortex by hippocampal signals. BioRxiv.

Fürth D, Vaissière T, Tzortzi O, Xuan Y, Märtin A, Lazaridis I, Spigolon G, Fisone G, Tomer R, Deisseroth K, Carlén M, Miller CA, Rumbaugh G, Meletis K (2018) An interactive framework for whole-brain maps at cellular resolution. Nat Neurosci 21:139–149.

Garrett ME, Nauhaus I, Marshel JH, Callaway EM (2014) Topography and areal organization of mouse visual cortex. J Neurosci 34:12587–12600.

Glickfeld LL, Andermann ML, Bonin V, Reid RC (2013) Cortico-cortical projections in mouse visual cortex are functionally target specific. Nat Neurosci 16:219–226.

Glickfeld LL, Olsen SR (2017) Higher-Order Areas of the Mouse Visual Cortex. Annu Rev Vis Sci 3:251–273.

Huh CYL, Peach JP, Bennett C, Vega RM, Hestrin S (2018) Feature-Specific Organization of Feedback Pathways in Mouse Visual Cortex. Curr Biol 28:114–120.e5.

Iurilli G, Ghezzi D, Olcese U, Lassi G, Nazzaro C, Tonini R, Tucci V, Benfenati F, Medini P (2012) Sound-driven synaptic inhibition in primary visual cortex. Neuron 73:814–828.

Jurjut O, Georgieva P, Busse L, Katzner S (2017) Learning Enhances Sensory Processing in Mouse V1 before Improving Behavior. J Neurosci 37:6460–6474.

Kalatsky VA, Stryker MP (2003) New paradigm for optical imaging: temporally encoded maps of intrinsic signal. Neuron 38:529–545.

Keller AJ, Roth MM, Scanziani M (2020) Feedback generates a second receptive field in neurons of the visual cortex. Nature 582:545–549.

Keller GB, Bonhoeffer T, Hübener M (2012) Sensorimotor mismatch signals in primary visual cortex of the behaving mouse. Neuron 74:809–815.

Kim EJ, Zhang Z, Huang L, Ito-Cole T, Jacobs MW, Juavinett AL, Senturk G, Hu M, Ku M, Ecker JR, Callaway EM (2020) Extraction of Distinct Neuronal Cell Types from within a Genetically Continuous Population. Neuron.

La Chioma A, Bonhoeffer T, Hübener M (2019) Area-Specific Mapping of Binocular Disparity across Mouse Visual Cortex. Curr Biol 29:2954–2960.e5.

Lee AM, Hoy JL, Bonci A, Wilbrecht L, Stryker MP, Niell CM (2014) Identification of a brainstem circuit regulating visual cortical state in parallel with locomotion. Neuron 83:455–466.

Leinweber M, Ward DR, Sobczak JM, Attinger A, Keller GB (2017) A sensorimotor circuit in mouse cortex for visual flow predictions. Neuron 95:1420–1432.e5.

Mao D, Neumann AR, Sun J, Bonin V, Mohajerani MH, McNaughton BL (2018) Hippocampus-dependent emergence of spatial sequence coding in retrosplenial cortex. Proc Natl Acad Sci USA 115:8015–8018.

Marques T, Nguyen J, Fioreze G, Petreanu L (2018) The functional organization of cortical feedback inputs to primary visual cortex. Nat Neurosci 21:757–764.

Marshel JH, Garrett ME, Nauhaus I, Callaway EM (2011) Functional specialization of seven mouse visual cortical areas. Neuron 72:1040–1054.

McGinley MJ, Vinck M, Reimer J, Batista-Brito R, Zagha E, Cadwell CR, Tolias AS, Cardin JA, McCormick DA (2015) Waking state: rapid variations modulate neural and behavioral responses. Neuron 87:1143–1161.

Minderer M, Brown KD, Harvey CD (2019) The Spatial Structure of Neural Encoding in Mouse Posterior Cortex during Navigation. Neuron 102:232–248.e11.

Murakami T, Matsui T, Ohki K (2017) Functional segregation and development of mouse higher visual areas. J Neurosci 37:9424–9437.

Nassi JJ, Cepko CL, Born RT, Beier KT (2015) Neuroanatomy goes viral! Front Neuroanat 9:80.

Niell CM, Stryker MP (2010) Modulation of visual responses by behavioral state in mouse visual cortex. Neuron 65:472–479.

Nitzan N, McKenzie S, Beed P, English DF, Oldani S, Tukker JJ, Buzsáki G, Schmitz D (2020) Propagation of hippocampal ripples to the neocortex by way of a subiculum-retrosplenial pathway. Nat Commun 11:1947.

Nurminen L, Merlin S, Bijanzadeh M, Federer F, Angelucci A (2018) Top-down feedback controls spatial summation and response amplitude in primate visual cortex. Nat Commun 9:2281.

Pakan JMP, Currie SP, Fischer L, Rochefort NL (2018) The impact of visual cues, reward, and motor feedback on the representation of behaviorally relevant spatial locations in primary visual cortex. Cell Rep 24:2521–2528.

Poort J, Khan AG, Pachitariu M, Nemri A, Orsolic I, Krupic J, Bauza M, Sahani M, Keller GB, Mrsic-Flogel TD, Hofer SB (2015) Learning Enhances Sensory and Multiple Non-sensory Representations in Primary Visual Cortex. Neuron 86:1478–1490.

Saleem AB (2020) Two stream hypothesis of visual processing for navigation in mouse. Curr Opin Neurobiol 64:70–78.

Saleem AB, Ayaz A, Jeffery KJ, Harris KD, Carandini M (2013) Integration of visual motion and locomotion in mouse visual cortex. Nat Neurosci 16:1864–1869.

Saleem AB, Diamanti EM, Fournier J, Harris KD, Carandini M (2018) Coherent encoding of subjective spatial position in visual cortex and hippocampus. Nature 562:124–127.

Shamash P, Carandini M, Harris KD, Steinmetz NA (2018) A tool for analyzing electrode tracks from slice histology. BioRxiv.

Sit KK, Goard MJ (2019) Distributed and retinotopically asymmetric processing of coherent motion in mouse visual cortex. BioRxiv.

Song JH, Choi W, Song Y-H, Kim J-H, Jeong D, Lee S-H, Paik S-B (2020) Precise mapping of single neurons by calibrated 3D reconstruction of brain slices reveals topographic projection in mouse visual cortex. Cell Rep 31:107682.

Speed A, Del Rosario J, Mikail N, Haider B (2020) Spatial attention enhances network, cellular and subthreshold responses in mouse visual cortex. Nat Commun 11:505.

Tyson A, Rousseau CV, Niedworok CJ, Margrie TW (2020) cellfinder: automated 3D cell detection and registration of whole-brain images. Zenodo.

Vangeneugden J, van Beest EH, Cohen MX, Lorteije JAM, Mukherjee S, Kirchberger L, Montijn JS, Thamizharasu P, Camillo D, Levelt CN, Roelfsema PR, Self MW, Heimel JA (2019) Activity in lateral visual areas contributes to surround suppression in awake mouse V1. Curr Biol 29:4268–4275.e7.

Vinck M, Batista-Brito R, Knoblich U, Cardin JA (2015) Arousal and locomotion make distinct contributions to cortical activity patterns and visual encoding. Neuron 86:740–754.

Wang Q et al. (2020) The allen mouse brain common coordinate framework: A 3D reference atlas. Cell 181:936–953.e20.

Wang Q, Burkhalter A (2007) Area map of mouse visual cortex. J Comp Neurol 502:339–357.

Wang Q, Gao E, Burkhalter A (2011) Gateways of ventral and dorsal streams in mouse visual cortex. J Neurosci 31:1905–1918.

Wang Q, Sporns O, Burkhalter A (2012) Network analysis of corticocortical connections reveals ventral and dorsal processing streams in mouse visual cortex. J Neurosci 32:4386–4399.

Waters J, Lee E, Gaudreault N, Griffin F, Lecoq J, Slaughterbeck C, Sullivan D, Farrell C, Perkins J, Reid D, Feng D, Graddis N, Garrett M, Li Y, Long F, Mochizuki C, Roll K, Zhuang J, Thompson C (2019) Biological variation in the sizes, shapes and locations of visual cortical areas in the mouse. PLoS ONE 14:e0213924.

Witter MP, Doan TP, Jacobsen B, Nilssen ES, Ohara S (2017) Architecture of the Entorhinal Cortex A Review of Entorhinal Anatomy in Rodents with Some Comparative Notes. Front Syst Neurosci 11:46.

Zhuang J, Ng L, Williams D, Valley M, Li Y, Garrett M, Waters J (2017) An extended retinotopic map of mouse cortex. elife 6.

Zingg B, Hintiryan H, Gou L, Song MY, Bay M, Bienkowski MS, Foster NN, Yamashita S, Bowman I, Toga AW, Dong H-W (2014) Neural networks of the mouse neocortex. Cell 156:1096–1111.

